# Multi-Region Brain Organoid –Fusion Organoid with Cerebral, Endothelial and Mid-Hindbrain Components

**DOI:** 10.1101/2025.01.20.633788

**Authors:** Anannya Kshirsagar, Hayk Mnatsakanyan, Sai Kulkarni, John Guo, Kai Cheng, Oce Bohra, Ram Sagar, Vasiliki Mahairaki, Christian E Badr, Annie Kathuria

## Abstract

Brain organoid technology has revolutionized our ability to model human neurodevelopment in vitro. However, current techniques remain limited by their reliance on simplified endothelial cell populations rather than a complete endothelial system. We engineered Multi-Region Brain Organoids (MRBOs) that integrate cerebral, mid/hindbrain, and complex endothelial organoids into one structure. Different from the earlier approaches based on isolated HUVECs, our endothelial organoids contain diverse vascular cell types, including vascular progenitors, mature endothelial cells, pericytes, proliferating angiogenic cells and stromal cells. Our strategy employs a sequential modulation of key developmental pathways to generate individual organoids, followed by optimized fusion conditions that maintain regional identities while supporting cellular integration. Single-nucleus RNA sequencing shows that MRBOs develop discrete neural populations specific to each brain region alongside specialized endothelial populations that establish paracrine signalling networks. Integration analysis with human fetal brain data shows that MRBOs contribute to 80% of cellular clusters found in human fetal brain tissue (Carnegie stages 12-16), whereas CellChat analysis identifies 13 previously uncharacterized endothelial-neural signalling interactions. Notably, we uncover endothelial-derived factors that support the persistence of intermediate progenitor populations during hindbrain development, but not during the cerebral one thereby revealing a new role for complex endothelial populations in regional brain patterning. This platform enables matching multiple developmental regions, fully incorporating the endothelial nature at the same time providing opportunities for studying neurodevelopmental disorders in which neural-endothelial interactions are disrupted. Our engineered MRBO system establishes a foundation for investigating complex neurodevelopmental processes, providing an enabling context closer to physiological relevance.

## Introduction

Neurological diseases are some of the biggest challenges of the present world since they impact more than one billion people globally.^1^ Even more so in neurodevelopmental disorders, for instance, it is estimated that 317 million children^2^ throughout the world suffered from developmental disabilities in 2019, which is a heavy social and economic burden on families and health care systems. The recent brain assembloid platforms have been enhanced in a manner that makes it possible to create combinations of cortical organoids, midbrain, and hindbrain regions^3–8^, thus bringing them one step closer to constructing a model of human brain development. Nevertheless, there are still two major issues: (1) while researchers have incorporated isolated endothelial cells with cortical organoids, the incorporation of endothelial system with multiple brain regions at the same time is unexplored^9,10^ and (2) currently, isolated human umbilical vein endothelial cells (HUVECs) are used^11,12^ instead of endothelial organoids, which contain multiple cell types including, vascular progenitors, mature endothelial pericytes, proliferating angiogenic, and stromal cells that better represent the native neurovascular environment.

In human fetal development, endothelial and neural systems develop in concert as the two systems are known to be interconnected by several feedback loops that orchestrate regional patterning.^13–15^ It is important to consider these interactions to create models of neurodevelopmental disorders across various brain regions. However, the prior studies which included endothelial cells were mainly based on the vascularization of cortical organoids^16^ but neglected the role of endothelial-derived factors on the development of other brain regions at the same time.

To address these fundamental gaps, we have developed Multi-Region Brain Organoids (MRBOs), a platform that recapitulates key aspects of human fetal brain development by integrating cerebral, mid-hindbrain, and endothelial organoids into a cohesive construct. Our data demonstrates that these region-specific organoids maintain distinct transcriptional profiles that closely correlate with human fetal brain regional identity.

This is the new generation of brain organoids that is a great improvement to the modeling of brain development and the pathophysiology of the fetal brain. With three main regions of the brain and endothelial system, the MRBOs allow the analysis of highly complex human brain function over time. This platform is especially relevant for the analysis of the effects of environmental factors, genetics, and therapies on the neurodevelopment processes in different regions of the brain. Such analyses are crucial for understanding the mechanisms underlying neurodevelopmental disorders such as bipolar disorder, autism spectrum disorder, and schrizophrenia^17,18^, where regional disconnection is a key feature that adversely impacts individuals throughout their life.

## Results

### Generation of Multi Region Brain Organoid

We generated multi-region brain organoids (MRBOs) using three healthy control iPSC lines through a systematic protocol that integrates cerebral, endothelial, and mid/hindbrain organoids (Fig. 1a and b). Cerebral organoids were generated through established protocols^19,20^ using dual-SMAD inhibition for neural induction, followed by maturation in three-dimensional culture conditions to promote the development of cortical structures.

**Fig. 1:**
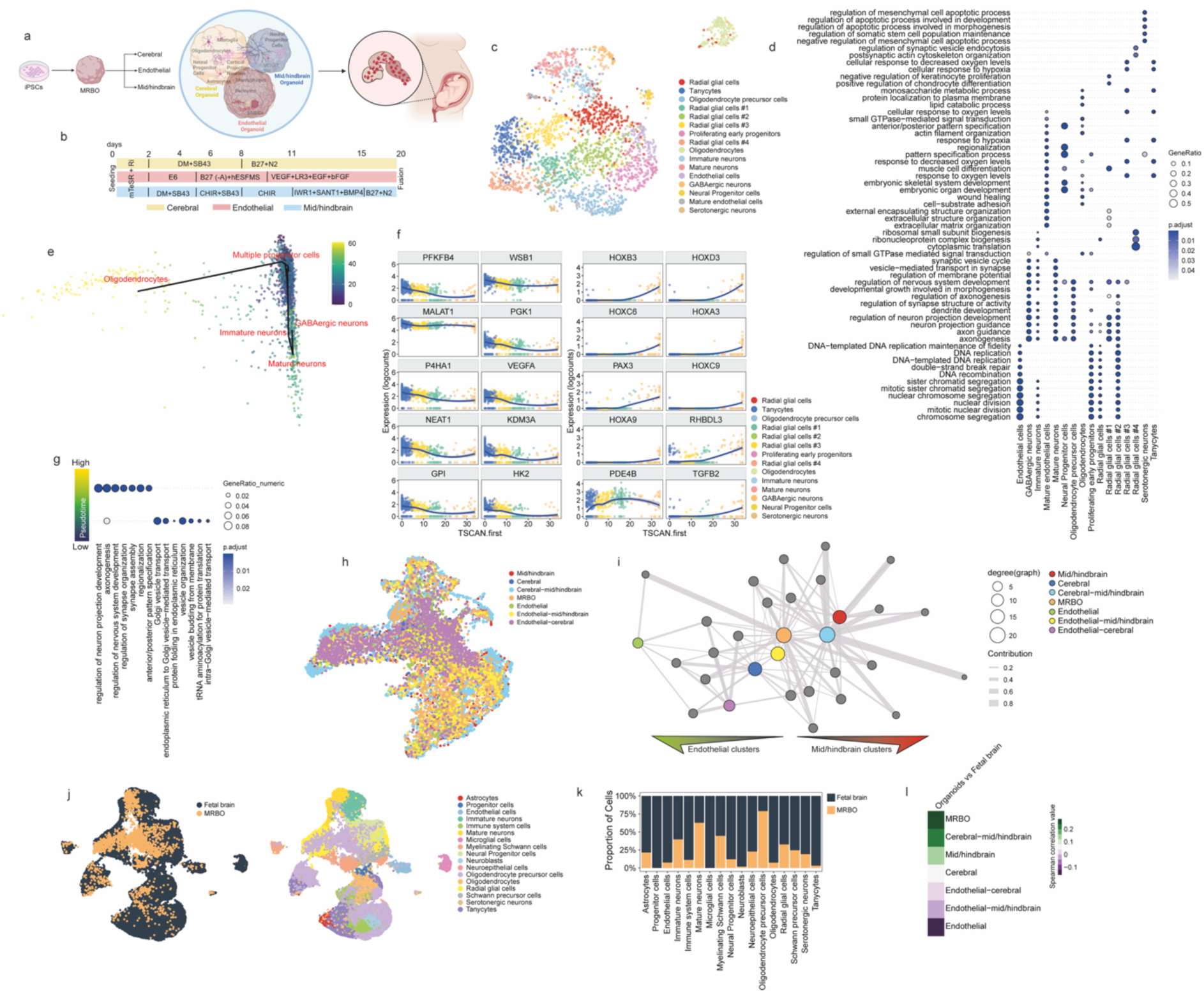
Generation and Molecular Characterization of Multi-lineage Brain Organoid Development Through Integrated Single-nucleus Analysis. (a) Basic principle and process used to form multi-region brain organoids (MRBOs) from iPSCs. (b) Timeline of protocol for generation of cerebral, endothelial, and mid/hindbrain organoids prior to fusion for MRBO formation. The cerebral organoids formation is governed by double SMAD inhibition, after day 5, the endothelial organoids are subjected to media containing retinoic acid and bFGF and all the mid/hindbrain organoid media contain bFGF. (c) UMAP visualization of single-cell RNA expression data from multi-region brain organoids (MRBO), with cells colored by cell type annotations.(d) Gene Ontology (GO) analysis highlighting the top seven most significant terms associated with gene expression markers defining each cell type in the MRBO.(e) PCA visualization of neural cells in the MRBO, colored by pseudotime progression as determined by TSCAN.(f) Dot plot of trajectory analysis for neural cells in the MRBO performed with TSCAN, showing the top 10 most highly expressed genes at low pseudotime (right) and the top 10 most highly expressed genes at high pseudotime (left).(g) GO analysis of the top 1,000 most highly expressed genes at high and low pseudotime in the neural component of the MRBO, represented as a dot plot.(h) UMAP visualization of the merged and integrated single-cell RNA expression data from MRBO, cortical organoids, endothelial organoids, mid/hindbrain organoids, cortical-endothelial organoids, mid/hindbrain-endothelial organoids, and cortical-mid/hindbrain organoids.(i) Interconnected network visualization showing the contributions of individual organoids to the integrated dataset and their relative contributions to each cell type cluster.(j) UMAP visualization of the integrated dataset combining MRBO and single-cell RNA sequencing data from the human fetal brain, colored by dataset origin (left) and clustered by gene expression profiles (right).(k) Bar plot depicting the relative contributions of each cluster from the fetal brain and MRBO.(l) Heatmap of Spearman correlation values representing gene expression similarities between each organoid type (assembloid or individual) and the fetal brain.

The endothelial organoids were developed in parallel from the same iPSC lines. Following embryoid body (EB) formation (days 0-2), cells were directed toward mesodermal lineage through WNT activation and BMP4 stimulation (days 2-9)^21^. These EBs were then embedded in a 1:1 collagen I and fibronectin matrix, with subsequent vascular fate specification achieved through VEGFA and FGF supplementation from day 10 onwards.

Mid/hindbrain organoids (MHOs) were generated using a modified protocol ^22,23^. While the initial five days followed standard neural induction, subsequent patterning was achieved using a specific combination of morphogens - CHIR, BMP4, IWR1, and SANT1. These organoids were embedded in Matrigel on day 15 and maintained in maturation medium supplemented with B27 and N2.

On day 20, we fused these three distinct organoid types using Matrigel as a supporting matrix. The successful integration was confirmed through light sheet microscopy (Supplementary Video 1). The resulting MRBOs were maintained until day 60 when comprehensive analyses were performed. This unified structure provides a robust platform for studying interregional interactions within a 3D brain microenvironment, enabling investigation of complex neurodevelopmental processes and disease mechanisms

To determine whether the multi-region brain organoid (MRBO) comprises all the cell types present in the individual organoids that form the assembly, we performed single-nucleus RNA sequencing analysis. The resulting UMAP visualization revealed distinct transcriptional clusters that captured the full cellular diversity of the MRBO (Fig. 1c). Through detailed annotation of these clusters based on canonical markers, we identified multiple neural populations alongside endothelial cell types, confirming the successful integration of both neural and vascular components. This comprehensive cellular map (Fig. 1c and S1a-d) demonstrated that MRBOs maintain the key cell populations from their constituent organoids while establishing a unified structure.

To better understand the gene expression signatures that define each cluster, we performed gene ontology (GO) analysis of the markers specific to each cluster (Fig. 1d). Cells representing advanced differentiation stages, such as neurons and oligodendrocytes, showed enrichment for GO terms related to neural development, axonogenesis, axon guidance, and forebrain development. In contrast, cells with stem-like features, such as early proliferating progenitors, radial glial cells, and oligodendrocyte progenitors, were enriched for terms related to cell proliferation and mitotic processes.

We further explored the developmental organization within the MRBO, we applied TSCAN to the single-nucleus RNA sequencing data to infer pseudotime and identify cellular trajectories in both neural and endothelial components of the MRBO. This analysis revealed insights into the spatial and temporal organization of the organoid, highlighting the distribution of cell types and their differentiation stages (Fig. 1e and S1e). High pseudotime values were observed in highly differentiated cells, such as mature neurons and oligodendrocytes, while low pseudotime values corresponded to cells with stemness properties and proliferative activity. Interestingly, the expression of genes upregulated along the pseudotime trajectory revealed that the HOX family of transcription factors was highly expressed in cells with high pseudotime (Fig. 1f and S1e), indicating the development of mid/hindbrain regions ^24^. Gene ontology analysis of the top genes expressed at low pseudotime confirmed enrichment for processes related to cell proliferation and mitosis, whereas genes expressed at high pseudotime were associated with neural development and morphogenesis (Fig. 1g). For the endothelial component, we observed a maturation trajectory along pseudotime, characterized by the expression of cell proliferation-related genes at low pseudotime, transitioning to genes associated with the synthesis of the endothelial extracellular matrix at higher pseudotime (Fig. S1f-i).

We evaluated the integration efficiency and cellular composition across different organoid combinations, we performed a comprehensive analysis by merging single-nucleus RNA sequencing data from all organoid types - individual components (cortical, endothelial, and mid/hindbrain organoids) and their various combinations (cortical-endothelial, mid/hindbrain-endothelial, and cortical-mid/hindbrain organoids), along with the complete MRBO. The resulting integrated UMAP visualization (Fig. 1h and S1j) revealed the global transcriptional landscape across all organoid configurations, enabling direct comparison of cellular populations and their distributions. This unified analysis approach allowed us to assess how cellular identities were maintained or modified across different organoid combinations, while providing insights into the successful integration of regional identities within the MRBO structure.

Dimensional reduction and clustering of the integrated object revealed an even distribution of MRBO-derived cells across all integrated datasets. The analysis of each organoid’s contribution to individual clusters revealed that the MRBO contributed to 20 out of 24 clusters (Fig. 1i), with contributions evenly distributed across endothelial, cerebral, and mid/hindbrain clusters, underscoring the robust integration of the component organoids. Additionally, we analyzed gene expression correlations among the different types of organoids (Fig. S1k) and found that the MRBO showed strong correlation with neural lineage-derived organoids (cerebral and mid/hindbrain), indicating that these are the most significant components of the assembly, although the endothelial component is also represented (Fig. S1l-n).

To assess the biological relevance of MRBO as an advanced model for brain development, we integrated MRBO data with single-nucleus RNA sequencing data from the human fetal brain. Dimensional reduction and clustering of the integrated dataset showed a substantial overlap between MRBO and the human fetal brain (Carnegie stage of human development 12-16)^25^ (Fig. 1k). Cells from the MRBO accounted for 85% of the total clusters after integration with the fetal brain organoid data (Fig. 1k) and emerged as the assembloid most closely correlating with the gene expression patterns of the fetal brain (Fig. 1l). These findings support MRBO as a relevant model for studying brain development and regional specification. In addition, we performed a reproducibility assessment on MRBOs. GAD65, a GABAergic neuron marker, showed excellent reproducibility (82.09%) between lines, while CTIP2, a deep layer neuron marker, maintained strong consistency (76.20%). The endothelial marker CD31 showed more variation between lines but maintained consistent expression patterns within each line. The mean overall reproducibility of 68.47% across all markers demonstrates the reliability of our organoid system for modeling neural development, with particularly strong performance in neuronal marker expression. (Supplementary table 1, staining not shown).

### Characterizing Cellular Heterogeneity Across Individual and Fused Brain Organoid Systems

To comprehensively characterize the cellular heterogeneity and validate the successful integration of distinct brain regions, we performed comparative single-nucleus RNA sequencing analysis across all organoid types - both individual components (cortical organoids, endothelial organoids, and mid/hindbrain organoids) and their various combinations (Fig.2). This systematic profiling approach allowed us to trace the contribution of each organoid type to the final assembled structure while confirming the preservation of region-specific cellular identities.

**Fig. 2:**
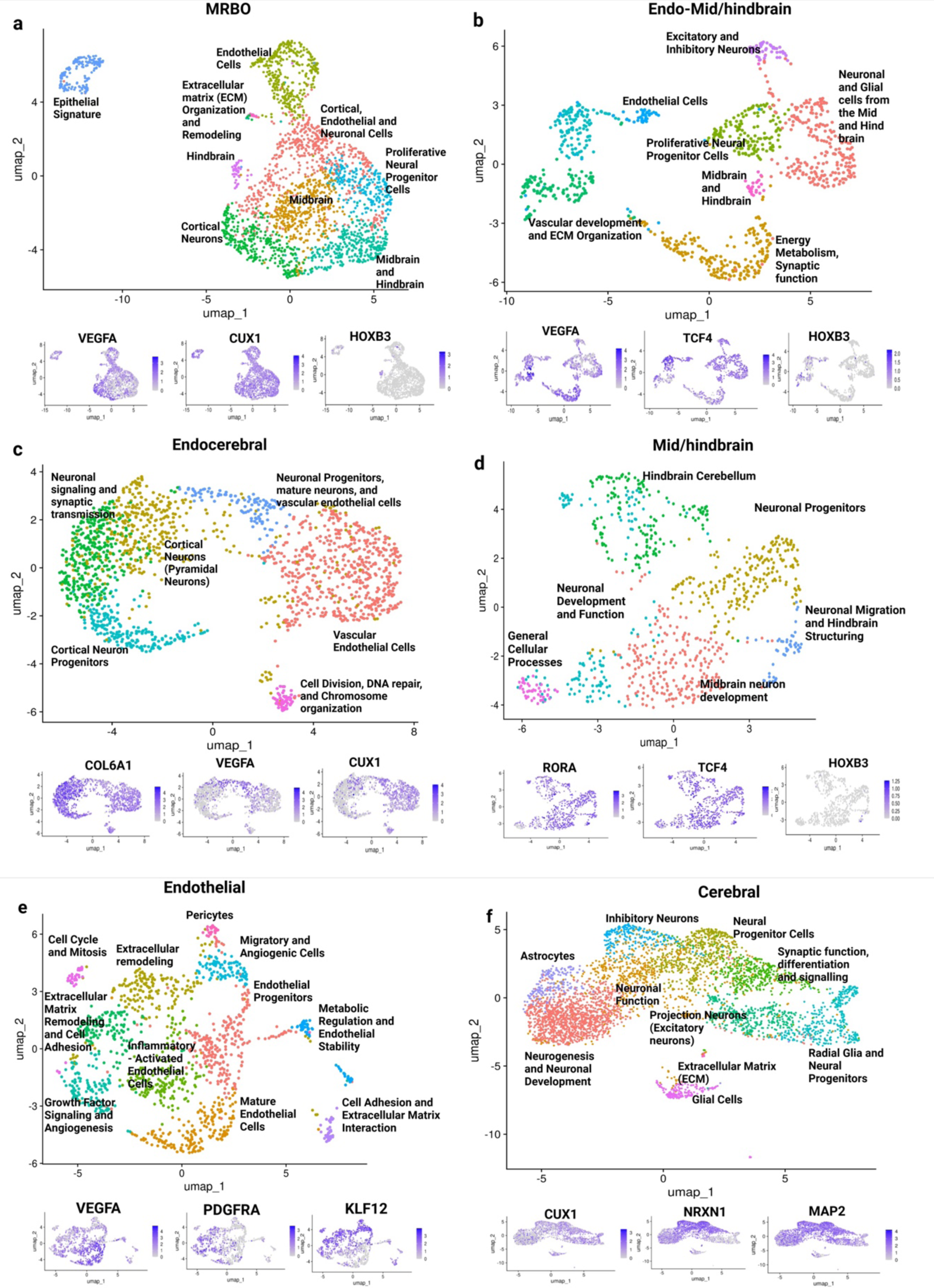
Transcriptomic characterization of MRBO, MHO, endothelial and cerebral organoid. **a)** MRBO snRNA-seq sample from 2 months of differentiation, showing expression of endothelial cell marker, VEGFA (left), neuronal differentiation marker, Cux1 and mid/hindbrain marker, HOXD3. **b)** The UMAP of Endo-Mid/brain depicts expression of endothelial markers like PDFGRA and mid/hindbrain neuronal markers – RORA and SOX5. **c)** UMAP of endocerebral show Cux1, COL6A1 and KLF12 gene expression signifying presence of both endothelial cells and prefrontal cortex neurons. **d)** UMAP of MHO show RORA, PBX1 and ROBO2 expression. **e) s**nRNA sequencing of endothelial organoid sample show expression of VEGFA, PDGFRA and KLF12; and that of **f)** cerebral organoid sample show expression of Cux1, NRXN1 and MAP2 markers. Single-nucleus RNA sequencing was performed using two independent lines (n=2 biological replicates per line). All organoids were analysed at day 60 of differentiation from iPSC stage.

We isolated and analyzed 5000 single nuclei from 2-month-old organoids of each type. To create an integrated view of cellular populations across all samples, we employed a comprehensive computational workflow. Post-sequencing data integration was performed using the Seurat pipeline, with dimensionality reduction achieved through Uniform Manifold Approximation and Projection (UMAP). This integrated analysis approach enabled us to generate detailed transcriptional maps that revealed how individual cell populations from each organoid component were maintained and combined within the final MRBO structure Graph-based clustering identified distinct cellular populations in the MRBOs and constituent organoids. In MRBOs, nine distinct clusters were visualized. Differential expression analysis highlighted the expression of key genes such as VEGFA (vascularization marker), CUX1 (cortical neuron marker), and HOXB3 (hindbrain marker), as shown in the overlay clusters (Fig. 2a). These findings confirm the presence of region-specific cellular identities within the MRBOs. Fusion-specific clusters were also analyzed. The endothelial and MHO fusion (Endo-MHO) organoids revealed seven distinct clusters with differentially expressed genes including VEGFA (vascularization marker), TCF4 (midbrain marker) and HOXB3 (hindbrain marker), which were localized to specific populations within the organoids (Fig. 2b). Similarly, six clusters were identified in Endocerebral organoids (Fig. 2c), with another six clusters observed in MHOs (Fig. 2d). Each cluster corresponded to unique cellular identities, indicative of successful differentiation and functional specialization. Further clustering analysis of endothelial organoids revealed eleven distinct clusters, comprising various cell types such as inflammatory cells, growth-factor signaling populations, angiogenesis-related cells, and mature endothelial cells (Fig. 2e). The diversity of cell types reflects the robust vascularization and cellular specialization achieved in the endothelial organoids. Cluster separations were assessed using silhouette width analysis, with all clusters showing distinct separations, confirming clear boundaries and minimal overlap between clusters. Canonical marker genes and highly differentially expressed genes were used to manually annotate each cluster with their corresponding cellular identities, further validating the specificity and accuracy of the differentiation and fusion protocols.

### Endothelial Cells Support Hindbrain Development Through Paracrine Signalling and Intermediate Progenitor Maintenance

Endothelial cells play a crucial role in the development and morphogenesis of the central nervous system. We hypothesized that endothelial cells play a crucial role in hindbrain development, as evidenced by the observed increase in HOXB3 expression correlating with elevated endothelial cell expression (Fig. 2a-b). ^26^ This phenomenon was not observed with TCF4 a midbrain marker (Fig. 2d). In addition, previous studies have established that HOX genes are integral to hindbrain formation, suggesting a potential regulatory interaction between endothelial cells and HOX gene expression.^27^ To test this hypothesis, we merged and integrated data from multi-region brain organoids (MRBO), endothelial organoids, and mid/hindbrain organoids, both individually and in various combinations (Fig. 3a-b).

**Fig. 3.**
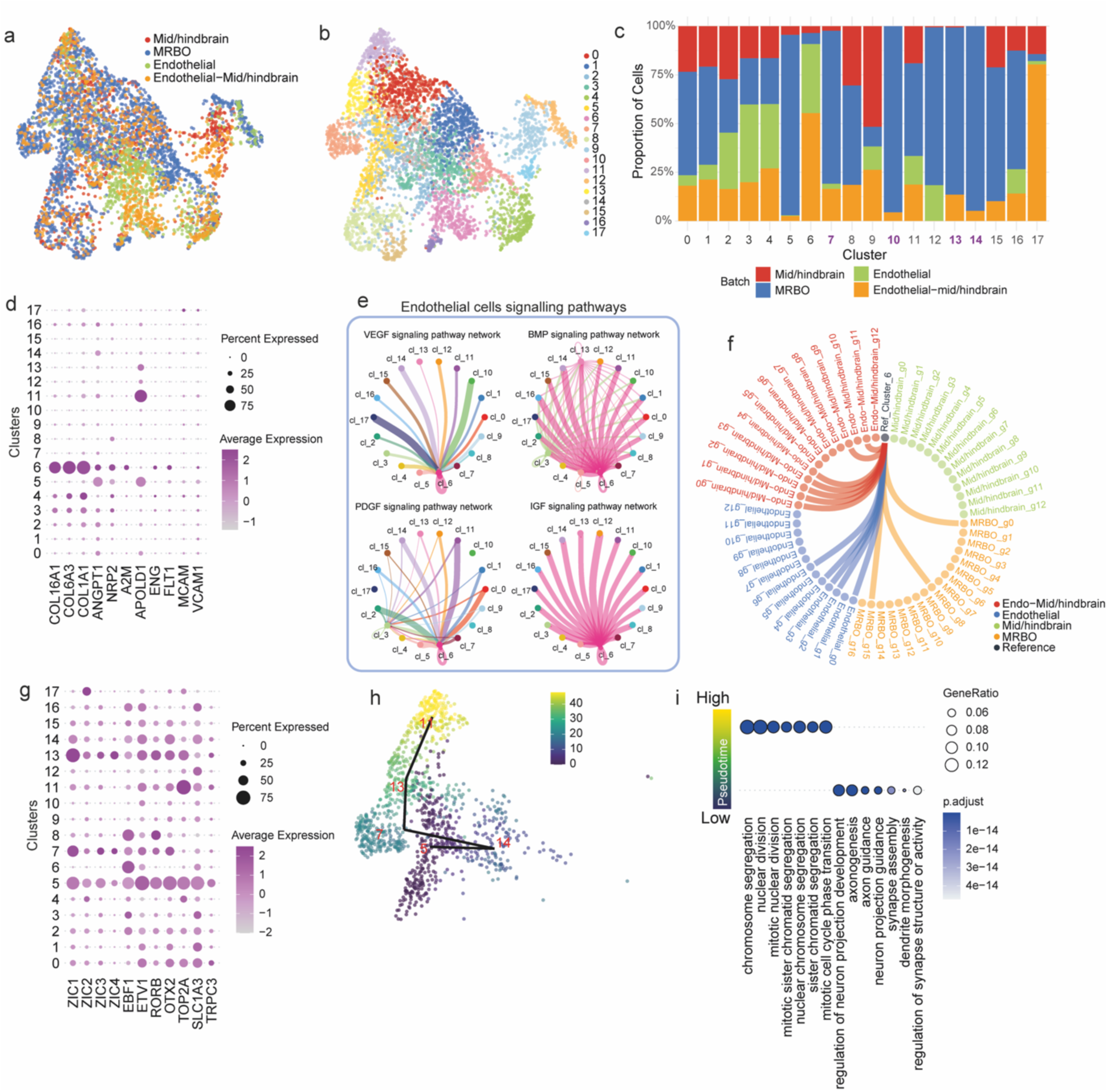
Analysis of Endothelial Cell Heterogeneity and Signaling Pathways. (a) UMAP visualization of the merged and integrated single-cell RNA sequencing data from multi-region brain organoids (MRBO), endothelial organoids, mid/hindbrain organoids, and endothelial-mid/hindbrain organoids, colored by batch origin.(b) UMAP visualization of the integrated dataset, colored by cell clusters.(c) Bar plot showing the relative contributions of each cell cluster from each batch origin.(d) Dot plot showing the expression of endothelial cell markers across cell clusters.(e) CellChat analysis illustrating the number of significant ligand-receptor pairs involved in endothelial signaling between clusters, with edge width proportional to the number of significant interactions.(f) Chord diagram depicting the Spearman correlation between cluster 6 of the integrated object and all clusters from each batch origin. Only correlations greater than 0.7 are displayed. (g) Dot plot highlighting the expression of hindbrain-specific markers across cell clusters in the integrated object.(h) UMAP visualization of cell clusters, colored by pseudotime as determined by TSCAN.(i) Gene Ontology (GO) analysis of the top 1,000 most highly expressed genes at high and low pseudotime, represented as a dot plot.Single-nucleus RNA sequencing was performed using two independent lines (n=2 biological replicates per line). All organoids were analysed at day 60 of differentiation from iPSC stage.

We performed clustering based on transcriptional profiles and analyzed the contribution of each individual organoid type to the resulting clusters (Fig. 3c). This analysis revealed that cluster 6 was uniquely present in organoids containing endothelial components, while it was absent in organoids made up solely of mid/hindbrain tissue. To confirm the endothelial identity of cluster 6, we examined the expressions of key endothelial cell markers, including FLT1, COL16A1, COL6A3, COL1A1, ANGPT1, and NRP2 (Fig. 3d). These markers were found to be highly expressed in cells within cluster 6, validating their endothelial nature. To evaluate the paracrine signaling contributions of these endothelial cells, we used CellChat to infer intercellular communication pathways among clusters, focusing on signaling pathways related to neural development and endothelial cells (Fig. 3e, Fig. S4a). Our analysis highlighted four key developmental signaling pathways: VEGF, BMP, PDGF, and IGF (Fig. 3e). ^28^ This cluster exhibited unique signaling behavior, including autocrine VEGF signaling and isolation to produce BMP and IGF factors. In addition, it is the prime known recipient of signals emanating from surrounding neural tissues in response to VEGF.

In the VEGF signaling pathway, cluster 6 demonstrated autocrine signaling and received VEGF signals from all other clusters. Notably, it was the only cluster expressing VEGF receptors. A similar trend was observed in the PDGF pathway, where both clusters 6 and 3 expressed receptors, although the signaling effect in cluster 3 was less prominent. These results suggest that cluster 6 cells function not only as endothelial cells but also as vascularization-supporting cells (Fig. 3f). In contrast, for the BMP and IGF pathways, cluster 6 was the exclusive producer of these cytokines, while the other clusters exclusively expressed receptors for these factors. This finding highlights the unique signaling role of cluster 6. It recapitulates early neurovascular interactions and molecular signatures of developing blood vessels. However, additional factors may be needed to achieve complete blood-brain barrier functionality.

To explore how endothelial cells influence hindbrain development, we examined the expression of hindbrain-specific markers. Five clusters (5, 7, 11, 13, and 14) were identified as expressing cerebellum-associated markers (Fig. 3g). Interestingly, clusters 7, 13, and 14 were only present in organoids containing both hindbrain and endothelial components. To investigate the developmental trajectories of these hindbrain-associated clusters, we isolated the cells in clusters 5, 7, 11, 13, and 14 and performed pseudotime analysis (Fig. 3h and S4b). This revealed a developmental progression beginning with cluster 5 (lowest pseudotime), followed by clusters 14, 7, 13, and finally 11 (highest pseudotime).

To understand the biological implications of this pseudotime progression, we performed gene ontology (GO) analysis on the top 1,000 genes expressed in cells with low pseudotime and the top 1,000 genes expressed in cells with high pseudotime (Fig. 3i). Cells with low pseudotime were enriched for GO terms related to axonogenesis and axon guidance, indicative of differentiated neural cells. In contrast, cells with high pseudotime were associated with terms related to cell proliferation, consistent with a progenitor cell identity. Interestingly, while progenitor cells (high pseudotime) and differentiated cells (low pseudotime) were present in all organoids containing mid/hindbrain components, intermediate progenitors (clusters 7, 13, and 14) were only found in organoids with endothelial components. This finding suggests that paracrine signaling from endothelial cells is essential for maintaining and proliferating intermediate progenitors during hindbrain development. Additionally, CellChat analysis further revealed thirteen distinct signaling networks mediating neurovascular communication, including WNT, NOTCH, and NCAM pathways (Fig. S4a).

We investigated whether the presence of the endothelial component influences the development of cerebral organoids. To do this, we merged and integrated all organoids containing a cerebral component, either alone or in combination with the endothelial organoid, as well as the endothelial organoid alone (Fig. S4c-e). Cells were clustered based on their gene expression signatures, and we analyzed each organoid’s contribution to the resulting clusters (Fig. S4f). For the cerebral component, we did not observe any significant influence from the presence of endothelial cells, concluding that endothelial cells primarily impact hindbrain development rather than cerebral one.

### Temporal Evolution of Neural Activity Patterns Across Different Organoid Types

Analysis of the neural activity patterns exposed distinct developmental trajectories for different organoid types over a 30-day period (Days 35-65). We found that in multi-region brain organoids (MRBOs) there was a significant increase in spike frequency (p < 0.05) and burst rate (p < 0.05) but a corresponding decrease in inter-spike interval (ISI) (Fig. 4a). found that in cerebral + midbrain/hindbrain organoids (Fig.4b), the developmental trajectory was more subtle. Although the changes in spike frequency and burst rate were not significant (ns), the ISI measurements did show consistent alterations over time. The spikes per average network burst, however, was significantly decreased (p < 0.01), indicating that the network activity is being refined. Significant increases in both spike frequency (p < 0.05) and burst rate (p < 0.01) were observed in mid/hindbrain organoids (Fig. 4c). The average network burst duration also showed a gradual increase, as did the mean ISI within the network bursts (p < 0.05), indicating the maturation of neural circuits. Cerebral organoids (Fig.4 d) showed different activity patterns; there was a peak in spike frequency at around Day 45-55 and then a slight decrease. The ISI, however, showed highly significant changes (p < 0.001), and the mean ISI within network bursts was significantly modulated (p < 0.05). All organoid types were analyzed for network normalized duration (IQR) (Fig.4e). The most significant changes were observed in the MRBOs. This suggests improvement in the network dynamics compared to cerebral and midbrain/hindbrain organoids over time.^29^ However, differences between individual organoid types were not statistically significant (ns).These findings, when taken together, show that various organoid types achieve distinct electrophysiological profiles over time, with MRBOs exhibiting the most dramatic shift in network activity patterns, which implies that different brain regions are adequately integrated for function.

**Fig. 4:**
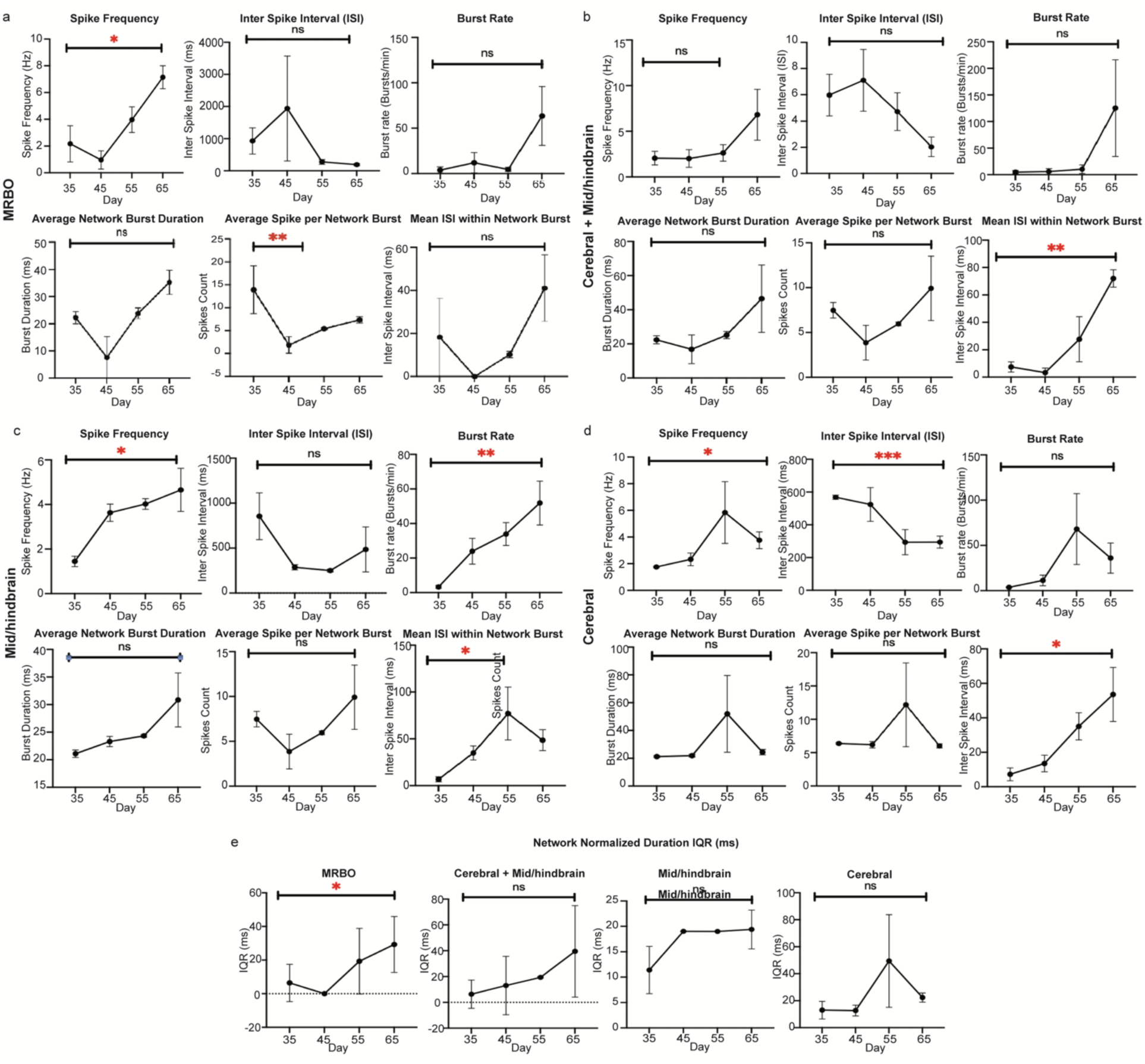
Network Maturation and Electrophysiological Development Across Distinct Brain Organoid Types: Electrophysiology data for the various organoids showing the Spike Frequency, Inter Spike Interval, Burst Rate, Average Network Burst Duration, Average Spikes Per Network Burst, Mean Inter Spike Interval Within the Network Burst. **a)** MRBO **b)** Cerebral + Mid/hindbrain Organoids **c)** Mid/hindbrain Organoids **d)** Cerebral Organoids. **e)** Network Normalized Duration IQR for MRBO, Cerebral + Mid/hindbrain, Mid/hindbrain, and cerebral organoids. Statistical analysis was carried out using the Dunnett’s multiple comparisons test and Welch’s t test where p-values show significance of the order: *P<0.05, **P<0.005, ***P<0.001. Experiment was performed using multiple iPSC control lines: MEA assay with three independent lines (n=3 biological replicates per line). All organoids were analysed at day 35 to 65 of differentiation from iPSC stage.

### Secondary Validation of Multi-Region Brain Organoids and characterization using immunohistochemistry

To further validate our single nuclear RNA sequencing (sc-RNA seq) on cellular composition and identity of our organoid systems, we used bulk RNA sequencing and immunohistochemistry. The bulk RNA sequencing analysis revealed transcriptional profiles that distinguish between cerebral, midbrain/hindbrain, and endothelial organoids and report expression of region-specific markers (Fig. 5a). The heatmap visualization of our data showed high expression of cortical markers (yellow cluster), mid/hindbrain markers (green cluster), and endothelial markers (brown cluster) in their corresponding organoid types, thus confirming correct regional specification. Quantitative analysis of key cellular markers in MRBOs found that there was a substantial presence of diverse cell types, with CD31, CTIP2, SV2A, VAChT, and CD34-positive cells present (Fig. 5b). This quantification showed that the fused organoid structure did contain multiple cellular lineages present and well maintained (additional quantification for other organoids is provided in supplementary Fig. 2). Immunohistochemical characterization of 2-month-old MRBOs showed strong expression of region-specific markers (Fig. 5c). The presence of PHOX2B confirmed hindbrain identity; CD34 validated the endothelial component; SV2A indicated mature synaptic development; and Cux1 and β-tubulin showed successful neuronal differentiation and maturation. Cerebral organoids^30^ had strong expression of myelin basic protein (MBP) and CUX1 marker for in layer II/III cortex (Fig. 5d). Nestin in sections confirmed that cortical development and maturation were properly recapitulated. Mid/hindbrain organoids (MHOs) also expressed specification multiple markers such as MAP2, TH, and PHOX2B (Fig. 5e). Angiogenesis and the beginning of new blood vessel formation were evident in the analysis of endothelial live cell imaging using Dil-Ac-LD^31^ (Fig. 5f). The two-month-old sections contained multiple endothelial and vascular markers, including CD34, VEGFR2, CD31, and PDGFβ (Fig. 5f). Thus, these results provide collective information and validation of cellular identities.

**Fig. 5:**
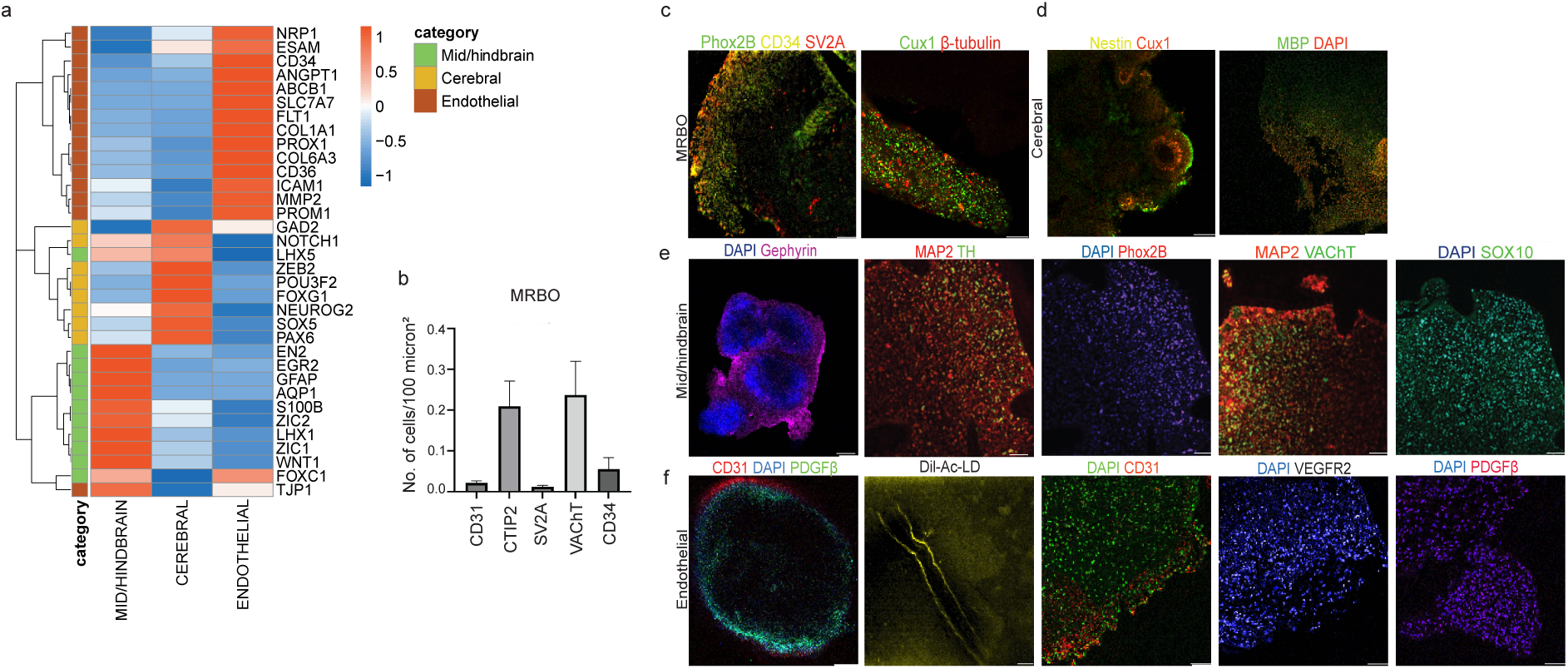
Validation of cell fate using immunohistochemistry analysis and bulk RNA sequencing. (a) Bulk RNA sequencing - Heatmap visualization of bulk RNA sequencing from cerebral organoids, midbrain/hindbrain organoids, and endothelial organoids, comparing the expression of cortical (yellow), hindbrain (green), and endothelial (brown) markers. (b) Number of cells expressing CD31, SV2A, VAChT and CD34 in multizone brain organoids per 100-micron square area. (c) Analysis of cell types in organoids using immunohistochemistry at 2 months, (top left to right) MRBOs show presence of PHOX2B, CD34, SV2A, Cux1 and β-tubulin. Scale bar 50µm. (d) Cerebral organoid showing presence of MBP, Nestin, Cux1, MBP, DAPI in sections at 2 months. Scale bar 50µm (left to right). (e) Immunostaining of whole mid/hindbrain organoid (MHO) (left to right) shows development of DAPI, Gephyrin, Sox 10, MAP2, TH, Phox2B. Scale bar 50µm. (f) Endothelial organoids depict presence of CD31 and PDGFβ (in whole organoid. Scale bar 150µm). The presence of angiogenesis is shown using Dil-Ac-LD dye validated by live cell imaging. Scale bar 25µm. Furthermore, markers such as CD34, VEGFR2, CD3, PDGFβ are observed in organoid sections at 1 month. Scale bar 50µm. (top left to right). Experiment was performed using multiple iPSC control lines: immunohistochemistry assay with three independent lines (n=3 biological replicates per line). All organoids were analysed at day 60 of differentiation from iPSC stage.

## Discussion

Our transcriptomic analysis reveals a remarkable similarity between Multi-Region Brain Organoids (MRBOs) and the human fetal brain, establishing this platform as a groundbreaking model for studying human brain development. The striking overlap of MRBO-derived cells with 80% of fetal brain clusters demonstrates these organoids’ capacity to recapitulate major neurodevelopmental events with unprecedented fidelity. This exceptional correlation with human fetal development positions MRBOs as an advanced experimental system for investigating complex neurodevelopmental disorders, particularly those affecting multiple brain regions. The incorporation of integrated endothelial components makes this platform especially valuable for studying conditions with vascular involvement, such as certain forms of autism and schizophrenia where blood-brain barrier alterations play a crucial role^13^.

A key discovery of our study is the identification of specific endothelial populations that actively support regional brain development through sophisticated paracrine signaling networks. The endothelial cluster (cluster 6), characterized by expression of classic endothelial markers such as FLT1, COL16A1, and ANGPT1, orchestrates multiple developmental pathways through VEGF, BMP, PDGF, and IGF signaling. Notably, we found that endothelial cells are essential for maintaining intermediate progenitor populations specifically during hindbrain development, as evidenced by the presence of intermediate progenitor clusters (clusters 7, 13, and 14, expressing hindbrain-specific markers) exclusively in organoids containing both hindbrain and endothelial components. This region-specific effect is particularly fascinating, as cerebral organoids showed significantly less dependence on endothelial influence. These findings not only advance our understanding of neurovascular development but also suggest new therapeutic strategies for disorders affecting mid/hindbrain development and function.

The electrophysiological characterization of MRBOs reveals another significant advancement: the development of functionally integrated neural networks that surpass the complexity observed in individual region organoids. Our analysis demonstrates progressive maturation of neural circuits, with significant changes in spike frequency, burst rates, and network synchronization patterns. This functional integration across different brain regions creates unprecedented opportunities for studying circuit development and dysfunction in a more physiologically relevant context. Such capabilities make MRBOs particularly valuable for investigating conditions characterized by altered brain connectivity and endothelial involvement, including cerebral cavernous malformations (CCM)^32^, CADASIL^33^, and vascular dementia^34^.

Despite these advances, we acknowledge certain limitations of our model. While MRBOs demonstrate excellent regional specification and cellular diversity, they currently lack a complete vascular network and long-range axonal projections. The absence of immune components and a fully functional blood-brain barrier presents opportunities for future improvements. However, recent work by Boutom et al. (2024)^13^ suggests that even non-perfused endothelial niches significantly enhance organoid fidelity and physiological relevance.

While extending culture periods could provide valuable insights into later developmental stages and circuit formation, the current model already offers substantial advantages for translational applications. Future iterations could further enhance physiological relevance through incorporation of additional cell types, such as microglia and other immune cells. Beyond their value in basic research, MRBOs represent a transformative platform for personalized medicine and drug development. By generating patient-specific MRBOs from induced pluripotent stem cells, this system enables precise modeling of individual disease phenotypes and personalized drug response testing. The presence of multiple brain regions alongside integrated endothelial components provides a more comprehensive testing environment than traditional 2D cultures or simpler organoid models. This enhanced physiological relevance could significantly improve the predictive power of preclinical drug screens, potentially reducing the high attrition rate currently seen in neuroscience drug development. Furthermore, the ability to systematically evaluate drug effects on both neural and vascular components within the same model offers unique opportunities for developing targeted therapeutics for complex neurovascular disorders. As we move toward more personalized treatment approaches, MRBOs could serve as a valuable tool in tailoring therapeutic strategies to individual patients, particularly for conditions affecting multiple brain regions or involving neurovascular interactions^28^. This advancement in organoid technology thus bridges a critical gap between traditional in vitro models and human clinical trials, offering a more reliable platform for therapeutic development in neurological and psychiatric disorders.

## Materials and Methods

### Human Induced Pluripotent Stem Cell Culture

We used three human induced-pluripotent stem cells (Johns Hopkins University) that were cultured in T75 flasks in Essential 8 medium containing DMEM/F12, L-glutamine and sodium bicarbonate (Gibco) at 1.734g/L on vitronectin. The cultures were maintained in T25 flask and passaged at 1:2 ratio in T75 flask after 5-7 d by versene (Gibco) for 4 min at 37°C and mechanical scrapping using cell scrappers. All the human pluripotent stem cells were maintained below passage number 50, and we also tested to confirm negative for mycoplasma presence. The three healthy control lines are all female, NIBSC8 (15-week-old) ^36^, WT4 (56-year-old) ^37^ and EP1NRB3 (16-week-old). ^38^

### Generation of Cerebral Organoids

The cerebral organoids are generated using control human induced-pluripotent stem cells (iPSCs) over 14 days. This protocol requires daily media changes. To begin, iPSCs are washed with 1X HBSS (Gibco) and dissociated to single cells using Accutase. The single cells are seeded at 25,000 cells/well in ultra-low attachment 24-well Aggrewell plates (800mm, Stem Cell Technologies) to form embryoid bodies (EBs) in Essential 8 medium supplemented with DMEM/F12, L-glutamine, sodium bicarbonate (Gibco), and ROCK inhibitor Y-27632 (10µM, 1:1000) for the first 24 hours. After 24 hours, the medium is changed to neural induction medium containing dual SMAD inhibitors: 10µM SB431542 (TGFβ inhibitor) and 0.1µM LDN193189 (BMP inhibitor) in Neural Basal Medium supplemented with N2 and B27. The EBs are maintained in this condition for 5-7 days to promote neuroectoderm formation. Following neural induction, the medium is changed to neural differentiation medium containing Neural Basal Medium with N2 and B27 supplements, without SMAD inhibitors. The EBs are maintained in this medium for 4-5 days to allow for neural progenitor expansion. On day 14, the neural spheroids are embedded in Matrigel droplets to provide three-dimensional scaffold support. The embedded organoids are then maintained in cerebral organoid maturation medium (Catalog # 08571). This medium supports the development of various neural cell types and the formation of organized neural tissues.

### Generation of Endothelial Organoids

The endothelial organoids are generated using control human induced-pluripotent stem cells in 10 days adapted from the Wimmer, R. (2019)^21^ protocol. The modifications are the use of hang drop with extra cell matrix Collagen (ThermoFisher Catalog #: 33016015 and Fibronectin (Gibco Catalog number A1048301-1:1) and induce more endothelial production the use of Endothelial Cell Growth Media (R&D systems Catalog #: CCM027). First, the iPSCs were washed using 1X HBSS (Gibco) and dissociated with Accutase (Gibco) to generate single cells. These single cells were seeded at 25,000 cells/well concentration in ultra-low attachment 96-well plate (Nunc) to form embryoid bodies (EBs) in Essential 8 medium containing DMEM/F12, L-glutamine and sodium bicarbonate (Gibco) and ROCK inhibitor Y-27632 (10µM, 1:1000). EBs were maintained in this condition for 48 hr. On day 3, the medium was changed to mesoderm induction media consisting of DMEM/F12 supplemented with 1X B27 minus insulin (Gibco), 1X N2 (Gibco), 8 ng/mL Activin A (to activate TGF-β signaling), 5 ng/mL BMP4 (for mesoderm specification), 1 µM CHIR99021 (GSK3 inhibitor to activate WNT signaling), and 10 ng/mL FGF2 (to promote mesoderm differentiation). The EBs were maintained in this medium for the next two days to stimulate mesoderm fate commitment. Day 5-8, the EBs were subjected to vascular induction medium allows the organoids to form appropriate vasculature network. On day 9, the EBs were embedded in Collagen and Fibronectin (1:1) and fed with Endothelial Cell Growth Media (R&D systems Catalog #: CCM027).

### Generation of Mid/hindbrain Organoids (MHO)

The hind and midbrain organoids are generated using control human induced-pluripotent stem cells in 14 days (about 2 weeks). This protocol required media changes every day. First, the iPSCs were washed using 1X HBSS (Gibco) and dissociated with Accutase (dissociative enzyme) to generate single cells. These single cells were seeded at 25,000 cells/well concentration in ultra-low attachment 24-well Aggrewell plate (Stem cell Technologies) to form embryoid bodies (EBs) in Essential 8 medium containing DMEM/F12, L-glutamine and sodium bicarbonate (Gibco) and ROCK inhibitor which is a Rho kinase inhibitor; Y-27632 (10µM, 1:1000). EBs were maintained in this condition for 24 hr. and then medium was changed to MHO media 1 (Gibco) that containing 2µM dorsomorphin (to inhibit BMP signaling, batch 233309), 10 µM SB431542 (TGFβ pathway inhibitor, batch 262852), 10ng/mL bFGF (fibroblast growth factor). The EBs were maintained in this medium for 48 hrs and then transferred to MHO media 2 containing 3µM CHIR99021 (to upregulate Wnt/β-catenin signaling pathway, batch 232931), 10µM SB431542 and, 10ng/mL bFGF for 3 days. After 3 days, the media was changed to MHO media 3 containing 3 µMCHIR99021 and 10ng/mL bFGF. The EBs were maintained in this media for the next 3 days. Finally, the media was changed to MHO media 4 containing 10µM IWR1 (block Wnt/β-catenin signaling, batch 280452), 250nM SANT1 (inhibits Sonic Hedgehog (Shh) signaling pathway, batch 157339), 25ng/mL BMP4 (Bone morphogenetic protein 4 to stimulate neural stem cell differentiation, batch 252798) and 10ng/mL bFGF for next 4 days. On day 15, these EBs were embedded in Matrigel. The organoids were further maintained in MHO maturation media containing NeuroBasal media (Gibco) with B27 (Gibco) and N2 supplement (Gibco) (supplements to improve neuronal differentiation).

### Generation of MRBOs

The multi-region brain organoids were formed by fusion of cerebral organoids, endothelial organoids and mid/hindbrain organoids using the hang-drop method. Each organoid type was positioned in proximity and coated with Matrigel (CLS356230) to facilitate adhesion and fusion. Using the Agreewell system each organoid before fusion was 800μm, after fusion the organoid, reached the final size ∼2-3 mm. An observation was made that endothelial organoid absorbed the cerebral+mid/hindbrain organoid (Supplementary video 1). Media compostion was a 1:1:1 mixture of maturation media (Neural/Endothelial/Brain Region), this combined cerebral organoid maturation media (STEMCELL Technologies, Catalog # 08571), endothelial organoid maturation media (R&D systems Catalog #: CCM027), and mid/hindbrain organoid (MHO) maturation media (see above).

### Immunohistochemistry

The MRBOs were fixed with 4% paraformaldehyde (PFA) for 1 h, washed with 1X PBS (Gibco) and incubated in 30% sucrose overnight at 4°C. The sucrose was removed by 3 PBS washes for 10 min each. The organoids were then paraffin embedded and sectioned to get sections on a slide. After which, the slides were rinsed in 1X PBS (Gibco) and permeabilized with 1% Triton X-100 in PBS for 10 min. The sections were blocked using blocking solution (10% goat serum in PBS) for 1 h at room temperature. Primary antibodies were diluted in antibody solution (1% goat serum in PBS) and the samples were incubated in the antibody overnight (18 h). Samples were then washed by giving washes with two PBST (1% Triton X-100) washes for 10 min each. The samples were given a single PBS wash for 10 min before incubating them in secondary antibodies (made in antibody solution) for 1 h in the dark. Rinse out the secondary antibodies with PBST and PBS washes, 10 min each. Sections were stained with Hoechst (Invitrogen) and then using mounting media (Anti-Fade Prolong Gold, Invitrogen) coverslip the sections. The slides were imaged on the IXM High Content Imager or the Leica SP8 microscope.^39,40^ The live cell imaging of endothelial organoid is done using Dil-Ac-LDL uptake kit (Cell applications inc., lot number - 022). The light sheet microscopy was performed by clearing the MRBO sample using RapiClear for a week followed by antibody staining. The sample was imaged using Olympus UltraMicroscope.^41^

### snRNA - Sequencing Analysis

RNA integrity was assessed using an Agilent 4200 Tapestation, ensuring RNA Integrity Number (RIN) values above 5.0 for all samples. A total of 150 ng of RNA per sample was used as input for library preparation. Libraries were prepared using the NEBNext UltraExpress RNA Library Prep Kit (New England Biolabs, Ipswich, MA), following the manufacturer’s protocol. mRNA was enriched using poly(A) selection. Enriched mRNA was fragmented, reverse transcribed, and converted to cDNA. cDNA underwent end repair, adapter ligation, and size selection. PCR amplification was performed using 11 cycles to ensure sufficient library yield while minimizing amplification bias. Library quality and size distribution were assessed using a Tapestation (Agilent) and quantified by Qubit (Thermo Fisher Scientific). Final libraries were pooled equimolarly and sequenced on an Illumina NovaSeq X Plus platform using a 25B lane configuration with paired-end 150 bp (PE150) reads with a minimum of 20M Reads per sample. The sequencing run achieved an average depth of 20M reads per sample. Base calling and demultiplexing were performed using Illumina’s DRAGEN Bio-IT platform. Raw sequencing reads were processed for quality control using FastQC and trimmed to remove adapters and low-quality bases using Trim Galore. Cleaned reads were aligned to the reference genome GRCh38.p14 using STAR. Quantification of gene expression levels was performed using Salmon. Final, statistical analyses were performed using DESeq to identify differentially expressed genes. All scripts and pipelines are available upon request to ensure reproducibility.

### Single-nucleus RNA Sequencing Data Processing

The single-nucleus RNA sequencing (scRNA-seq) data were analyzed using the Seurat package (version 5.0) in R. Quality control measures were applied to filter low-quality cells, and cells were clustered based on their gene expression profiles. Dimensionality reduction was performed using Uniform Manifold Approximation and Projection (UMAP) to visualize cellular heterogeneity. Cell type annotations were assigned by identifying cluster-specific marker genes and comparing them to established brain cell type signatures, facilitated by the sc-type package.

### Pseudotime Analysis

To investigate cellular trajectories and dynamic processes, pseudotime analysis was conducted using TSCAN^42^. Pseudotime values were inferred for each cell, and the trajectories were visualized on a UMAP embedding to highlight the progression of cells along developmental or differentiation pathways.

### Gene Ontology Enrichment Analysis

Gene ontology (GO) enrichment analysis was performed using the enrichGO function from the clusterProfiler package.^43^. Differentially expressed genes identified from specific clusters or pseudotime states were subjected to GO analysis to determine enriched biological processes, molecular functions, and cellular components.

### Batch Integration

To integrate scRNA-seq data from different experimental batches, Seurat’s IntegrateLayers function was employed, utilizing the canonical correlation analysis (CCA) method^44^. This approach enabled the identification of shared features across batches while mitigating batch effects, ensuring the generation of a unified and biologically meaningful dataset.

### Cell-cell Communication Analysis

Intercellular communication between cell clusters was analyzed using the CellChat package^45^. Ligand-receptor interactions were inferred to identify signaling pathways and construct communication networks between cell populations. Interaction networks were visualized to elucidate the relationships between clusters and their potential signaling roles.

### Data Acquisition and Preprocessing

Single-nucleus RNA sequencing (snRNA-seq) data were acquired from multiple experimental conditions, including Cerebral, Cerebral + Mid/hindbrain (MHO), MRBO, Endothelial, endothelial + Mid/hindbrain, and Endothelial + Cerebral, Mid/hindbrain (MHO). The raw data was provided in Matrix Market format, accompanied by feature and barcode files. Data were imported into R using the ReadMtx function from the Seurat package, generating sparse matrix objects to optimize memory usage. Each dataset was subsequently converted into a Seurat object to streamline downstream analyses.

### Quality Control and Normalization

Quality control steps were implemented to ensure the integrity of the data. Low-quality cells were filtered out based on standard scRNA-seq metrics. Gene expression data were normalized using the NormalizeData function, and highly variable features were identified with the FindVariableFeatures function to retain the most informative genes for further analyses.

### Dimensionality Reduction and Clustering

Dimensionality reduction was performed via principal component analysis (PCA) using the RunPCA function, focusing on the top principal components. Cell clustering was achieved through the FindNeighbors and FindClusters functions, with clustering resolutions ranging from 0.5 to 2, depending on the dataset, to delineate distinct cellular subpopulations.

### Visualization

Uniform Manifold Approximation and Projection (UMAP) was applied for two-dimensional visualization of the high-dimensional data using the RunUMAP function. Cluster identities were visualized with DimPlot, and gene expression patterns of interest were illustrated using FeaturePlot. Boxplots showing gene expression distributions across different samples were generated using the ggplot2 package to highlight key marker genes. Computational Environment: All analyses were conducted in R version 4.2.3 using the Seurat, and ggplot2 packages.

### Electrophysiology

Electrophysiological recordings were obtained from hind and midbrain organoid, cerebral organoids and MRBOs that derived from patient’s iPSC using MED64-Presto MEA plate. Setup on days 35, 45 and 55. First, the MEA plate was coated with laminin and poly-D-lysine and incubated overnight. The organoids were stuck onto the MEA plate with a drop Matrigel using the hang drop method. This created contact between organoids and electrodes for accurate reading. Data preprocessing comprised multiple steps to ensure high-quality spike train data for analysis. First, a bandpass filter was applied to the raw electrophysiological data, restricting the signal to a frequency range of 0.5 to 3000 Hz. This was achieved using a fourth-order Butterworth filter, chosen for its flat frequency response within the passband, minimizing distortion of relevant neural signals. The 0.5 to 3000 Hz range was selected based on the typical frequency characteristics of neural activity. Following bandpass filtering, a notch filter centered at 60 Hz with a 2 Hz stopband was applied to eliminate power line noise, a common artifact in electrophysiological recordings. Additional notch filtering was performed at 60 Hz harmonics (120 Hz, 180 Hz) to further suppress residual power line interference.

### Statistical Analysis and details

Experimental validation was performed using multiple iPSC control lines: immunohistochemistry and MEA analyses were conducted with three independent lines (n=3 biological replicates per line), while single-nucleus RNA sequencing and bulk RNA sequencing were performed using two independent lines (n=2 biological replicates per line). All statistical analyses were performed using R (version 4.2.3) and GraphPad Prism (version 9.0), with data presented as mean ± standard error of the mean (SEM) unless otherwise specified. Single-nucleus RNA sequencing analysis (n=2 biological replicates each for each treatment) was performed with differential expression analysis using DESeq2 (false discovery rate [FDR] < 0.05, absolute log2 fold change > 1); multiple testing corrections were performed using the Benjamini-Hochberg method, while cluster stability was validated using bootstrap resampling (1000 iterations). Gene Ontology enrichment analysis was performed using clusterProfiler (FDR < 0.05, Benjamini-Hochberg adjusted), while significance in CellChat path analysis was determined by permutation tests (1000 permutations, P < 0.05). MEA recordings (n=3 biological replicates per condition) were analyzed using a two-way ANOVA, with Dunnett’s multiple comparisons test for between-group comparisons and repeated measures ANOVA with Tukey as a post-hoc test for temporal analyses within groups, applying Welch’s correction when Levene’s test indicated unequal variances. Number-based quantification (n=3 biological replicates, minimum 5 random fields per replicate) was normalized to either DAPI-positive nuclei or area (per 100 μm²) and subjected to analysis using either one-way ANOVA with Tukey’s post-hoc test or two-tailed Student’s t-tests for paired comparisons, normality being evaluated using the Shapiro-Wilk test. Bulk RNA sequencing analysis (n=2 biological replicates for each treatment) was done using DESeq2 with similar significance criteria to single-nucleus sequencing, with incorporation of principal component analysis on regularized log-transformed data, and hierarchical clustering using Euclidean distance. Integration with human fetal brain data used ComBat-seq for batch effects correction, integration quality was assessed through silhouette analysis and correlations were conducted using Spearman’s rank correlation coefficient. Significance of the pseudotime trajectory was assessed using generalized additive models in TSCAN (FDR < 0.05), and all analyses were performed blinded to experimental conditions where applicable. The analyses used several R packages, including Seurat (v5.0), DESeq2 (v1.34.0), and clusterProfiler (v4.2.2); exact P-values are in source data, and custom scripts are available upon request. All analyses were declared significant at *P < 0.05, **P < 0.01, and ***P < 0.001.

## Data availability statement

The authors declare that the data that support the findings of this study are available from the corresponding author upon request. GEO accession number GSE288165, will be available after publication.

## Acknowledgment.

We thank Dr. Lena Smirnova and Dr. Donald J. Zack for the control iPSC lines provided to us. The manuscript’s grammar was improved using Claude 3 Sonnet (Anthropic, version 2024), an AI language model, while maintaining all scientific content and conclusions by the authors.

