## Supplementary file for "Multi-Region Brain Organoid –Fusion Organoid with Cerebral, Endothelial and Mid-Hindbrain Components"

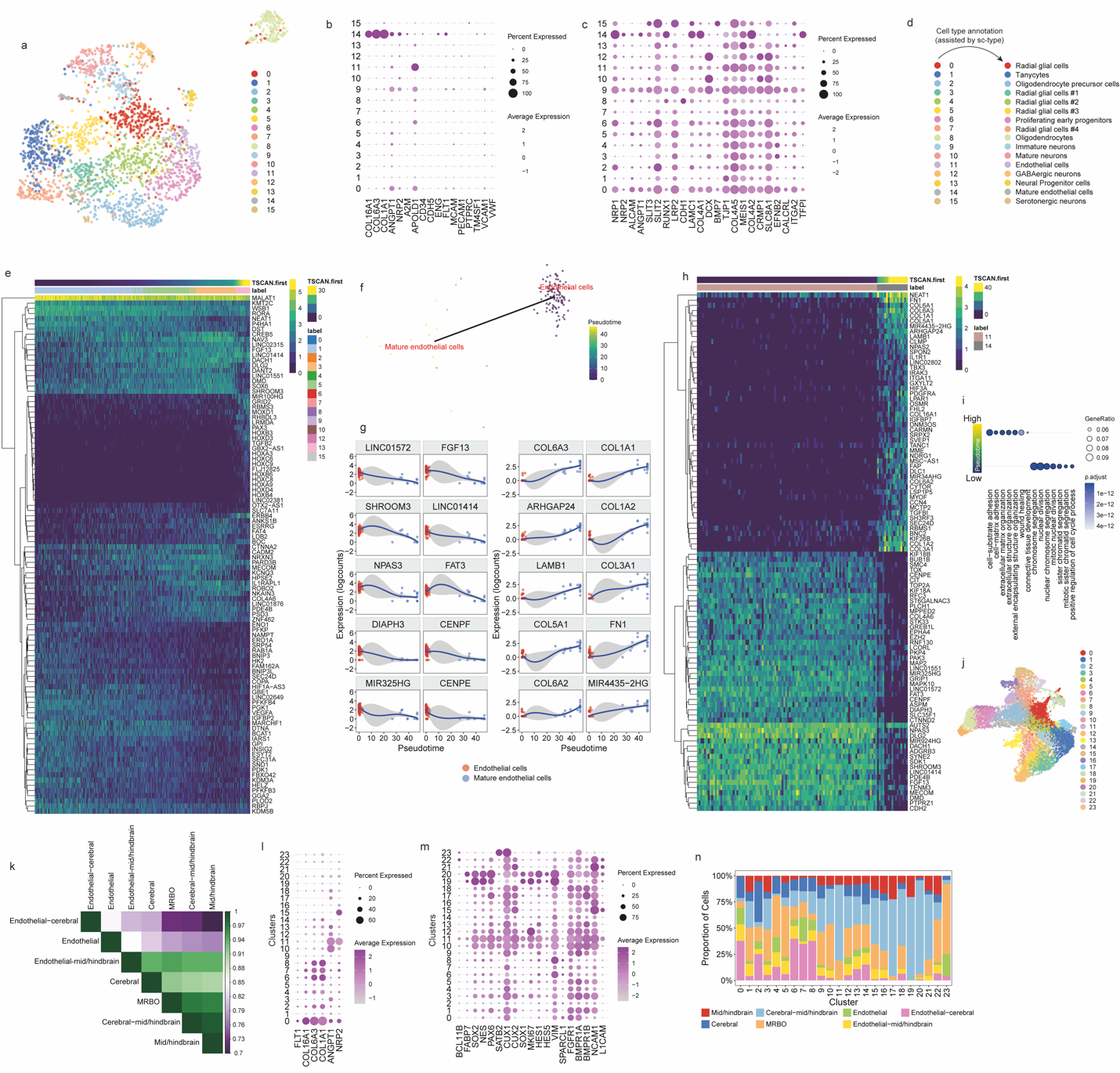

**Supplementary Figure 1:** (a) UMAP visualization of single-cell RNA expression data in MRBO, colored by gene expression clusters. (b) Dot plot showing the expression of endothelial markers across clusters in the MRBO.(c) Dot plot showing the expression of neural markers across clusters in the MRBO.(d) sc-type-assisted annotation of cell types associated with each cluster in the MRBO.(e) Heatmap of changes in gene expression during pseudotime progression in neural cells of the MRBO, analyzed using TSCAN.(f) PCA visualization of endothelial cells in the MRBO, colored by pseudotime progression as determined by TSCAN.(g) Dot plot of trajectory analysis for endothelial cells in the MRBO performed with TSCAN, showing the top 10 most highly expressed genes at low pseudotime (right) and the top 10 most highly expressed genes at high pseudotime (left).(h) Heatmap of changes in gene expression during pseudotime progression in endothelial cells of the MRBO, analyzed using TSCAN.(i) GO analysis of the top 1,000 most highly expressed genes at high and low pseudotime in the endothelial component of the MRBO, represented as a dot plot.(j) UMAP visualization of the merged and integrated single-cell RNA expression data from MRBO, cortical organoids, endothelial organoids, mid/hindbrain organoids, cortical-endothelial organoids, mid/hindbrain-endothelial organoids, and cortical-mid/hindbrain organoids, colored by clusters.(k) Heatmap of Spearman correlation values representing gene expression similarities among all generated organoids (assembloid or individual).(l) Dot plot showing the expression of endothelial markers across clusters in the integrated object resulting from merging MRBO, cortical organoids, endothelial organoids, mid/hindbrain organoids, cortical-endothelial organoids, mid/hindbrain-endothelial organoids, and cortical-mid/hindbrain organoids.(m) Dot plot showing the expression of neural markers across clusters in the integrated object resulting from merging MRBO, cortical organoids, endothelial organoids, mid/hindbrain organoids, cortical-endothelial organoids, mid/hindbrain-endothelial organoids, and cortical-mid/hindbrain organoids.(n) Bar plot illustrating the relative contributions of each cluster from all organoids constituting the merged and integrated object.

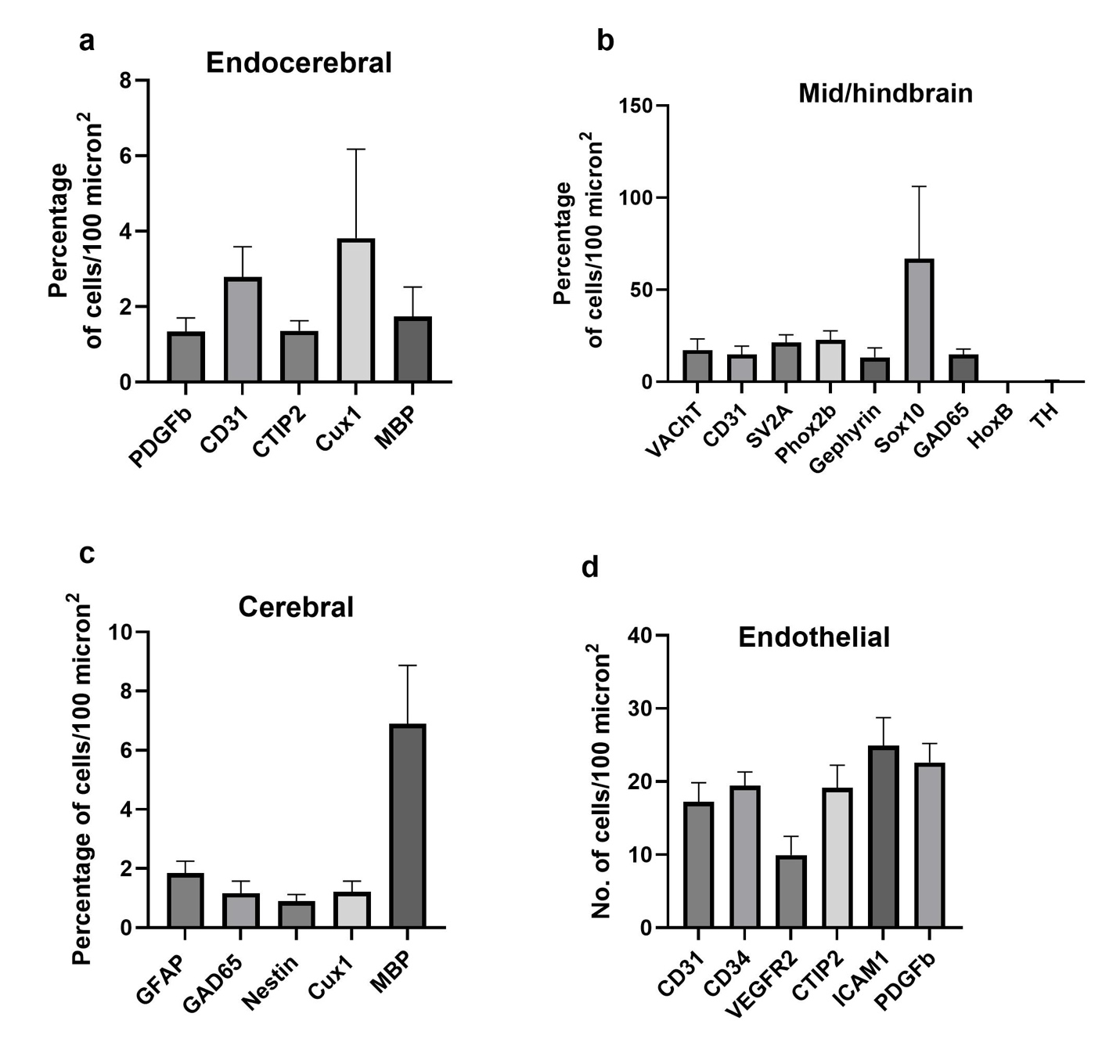

**Supplementary Figure 2: (a)** Percentage of cell types - PDGFβ, CD31 (endothelial cells), CTIP2 (layer V cortical marker), Cux1 (layer II/III cortical neurons), MBP (myelination), identified through immunofluorescence staining in MRBOs (n=5); **b)** VAChT (cholinergic neurons), CD31 (endothelial cells), SV2A (synaptic vesicle protein), Phox2b (autonomic neurons), Gephyrin (inhibitory synapses), Sox10 (neural crest cells), GAD65 (GABAergic neurons), HOXB (neural crest) and TH (dopaminergic neurons) were identified in Endocerebral organoids with SOX10 being present at highest percentage compared to the rest. **c)** The cerebral organoids contained higher percentages of IBA1 and MBP (myelin basic protein) followed by CD31 (endothelial cells) compared to other cell types – GFAP (astrocytes), GAD65 (GABAergic neurons), Nestin (neural progenitor cells) and Cux1 (layer II/III cortical neurons). **d)** Cell counting analysis of endothelial organoids identified cell types - CD31 (endothelial cells), CD34 (endothelial progenitor cells), VEGFR2 (Vascular endothelial growth factor receptor 2), CTIP2 (layer V cortical marker), ICAM1 (Intercellular adhesion molecule-1) and PDGFβ (platelet derived growth factor).

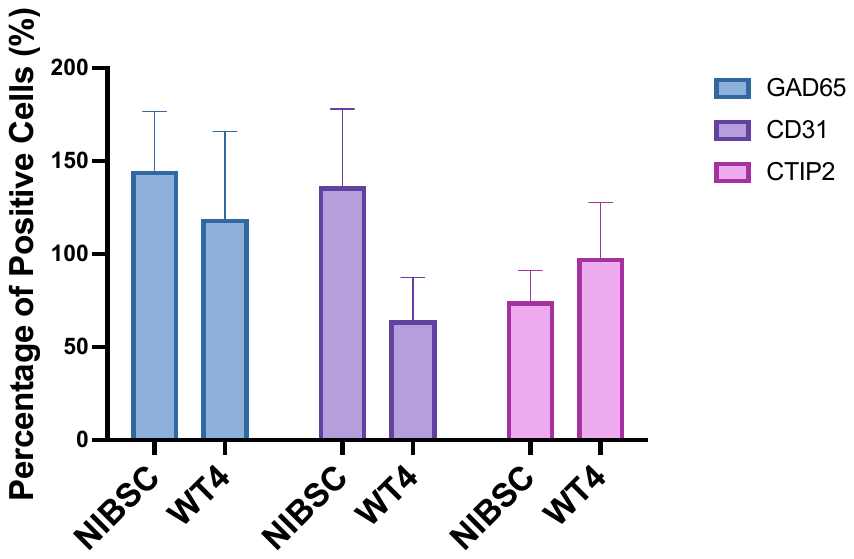

**Supplementary Figure 3:** Expression comparison of GAD65, CD31, and CTIP2 markers between NIBSC and WT4 cells. The graph shows the percentage of positive cells for each marker, with error bars representing standard deviation. Three distinct markers were analyzed: GAD65 (blue), CD31 (purple), and CTIP2 (pink).

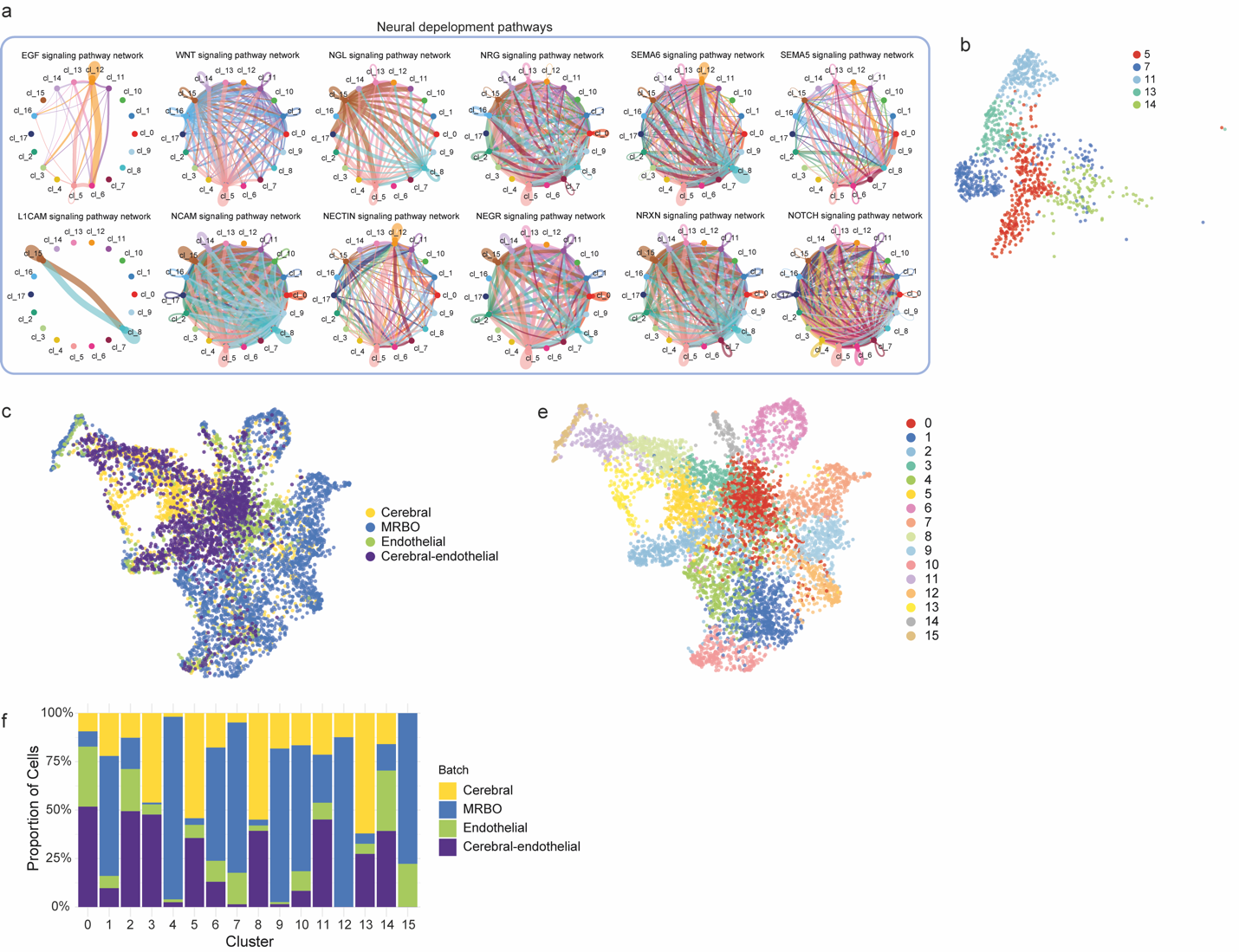

**Supplementary Figure 4**: (a) CellChat analysis illustrating the number of significant ligand-receptor pairs in signaling pathways related to neural development within the merged and integrated dataset of MRBO, endothelial organoids, mid/hindbrain organoids, and mid/hindbrain-endothelial organoids. Edge width is proportional to the number of significant interactions. (b) UMAP visualization highlighting clusters of interest (clusters 5, 7, 11, 13, and 14), colored by cluster identity. (c) Gene Ontology (GO) analysis showing the top seven most significant terms associated with gene expression markers defining the clusters of interest. (d) UMAP visualization of the merged and integrated single-cell RNA sequencing data from MRBO, endothelial organoids, cerebral organoids, and endothelial-cerebral organoids, colored by batch origin. (e) UMAP visualization of the integrated dataset, colored by cell clusters. (f) Bar plot illustrating the relative contributions of each cell cluster from each batch origin.

**Supplementary Table 1:**

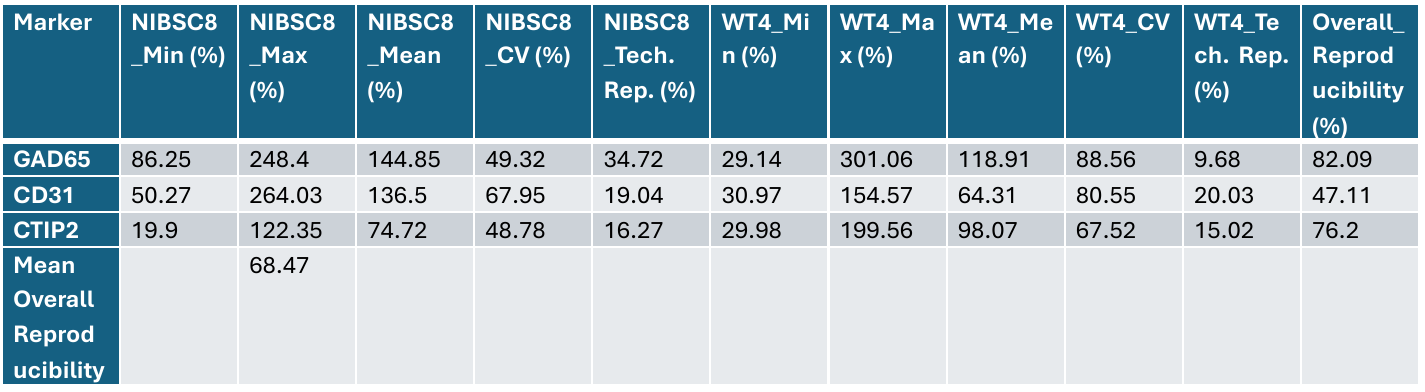

**Supplementary Method for Table 1 and Figure 3:**

**Comprehensive Reproducibility Analysis and Expression Levels by Cell Line**

Reproducibility and expression level analyses utilized immunocytochemistry data from two independent iPSC lines (NIBSC8 and WT4), with five technical replicates analyzed per marker for each line. For each section, the cell counts, and area measurements were obtained from immunostained sections, with areas measured in square micrometers (μm²). The percentage of positive cells was computed relative to total DAPI-positive nuclei. Expression levels for each marker (GAD65, CD31, and CTIP2) were quantified as the percentage of positive cells across the sections: Mean = (Σx)/n, where x represents individual percentage values and n is the total number of replicates; Eight defining metrics for technical reproducibility within each cell line; CV = (Standard Deviation/Mean) × 100Within-line technical reproducibility was calculated as the ratio of minimum to maximum value across the five technical replicates: Technical Reproducibility O = (Minimum Value/Maximum Value) × 100 Overall reproducibility between cell lines was assessed based on the comparison of mean expression values between NIBSC8 and WT4 lines: Overall Reproducibility = (Lower Mean/Higher Mean) × 100. The combined analysis provided both expression levels and reproducibility metrics. Expression data were plotted in bar graphs with error bars representing SEM. Overall reproducibility was presented with the expression data to provide a cumulative picture of marker consistency between lines. The mean overall reproducibility across all markers was calculated as the mean overall reproducibility among individual markers. Quality control measures instituted include area measurements standardized across samples, DAPI counterstaining as an internal control, and exclusion from analysis of sections yielding unsatisfactory staining quality or gross damage. Statistical analyses were performed with custom scripts coded in R (version 4.2.3).

**Light Sheet Microscopy:** Organoids were fixed in 4% paraformaldehyde (PFA) and subsequently immunostained for Phox2b (Rabbit anti-Phox2b, ThermoFisher Scientific #PA1-778), Beta-Tubulin (Mouse anti-Beta-Tubulin, Sigma-Aldrich #T8328), and CD31 (Rat anti-CD31, BD Biosciences #550274). Following antibody staining, the organoids were incubated overnight in RapiClear 1.47 clearing solution (SunJin Lab, #RC147002) at room temperature to achieve optical transparency. Cleared organoids were mounted in mineral oil (refractive index η = 1.47) and imaged using a light sheet fluorescence microscope (Ultramicroscope II, LaVision Biotec). The system employs a thin sheet of laser light to illuminate a single plane of the sample, minimizing photodamage and out-of-focus light. Fluorescence excitation was performed using lasers at 488 nm (Phox2b), 568 nm (Beta-Tubulin), and 785 nm (CD31). Serial optical sections were captured at a z-step of X µm, generating a stack of 257 high-resolution, 16-bit images for each stain. The resulting stacks were processed and reconstructed into 3D visualizations using ImageJ (3D Viewer plugin), enabling precise analysis of the spatial distribution of markers beneath the organoid surface.

**Supplementary Table 2:**

| **Antibodies** | **Dilution** | **Catalogue** |
| --- | --- | --- |
| Anti – SV2A | 1:200 | 119004 |
| Anti - VGLUT1 | 1:200 | 135011 |
| Anti - VAChT | 1:200 | 139103 |
| Anti - Gephyrin | 1:200 | 147111 |
| Anti - Homer1 | 1:200 | 160002 |
| Anti - GFAP | 1:200 | 173002 |
| Anti - MAP2 | 1:1000 | 188011 |
| Anti - GA2/GAD65 | 1:200 | 198104 |
| Anti - IBA1 | 1:200 | 234011 |
| Anti - MBP | 1:200 | 295002 |
| Anti - Nestin | 1:200 | 312111 |
| Anti - CD31 | 1:200 | HS-351004 |
| Hoechst (DAPI) | 1:2000 | H3570 |
| Anti - CTIP2 | 1:200 | ab18465 |
| Anti - PDGFβ | 1:200 | 3169T |
| Anti - VEGFR2 | 1:200 | MA5-15157 |
| Anti - ICAM1 | 1:200 | 60299-I-Ig |
| Anti - CD34 | 1:200 | 60180-I-Ig |
| Anti - Phox2B | 1:200 | 27870S |
| Anti - SOX10 | 1:200 | 89356T |
| Anti-TUBB3 | 1:1000 | 801213 |

**Supplementary Video**: Integration of region-specific organoids (cerebral, mid/hindbrain) with endothelial organoids to model complex brain development. CD31 endothelial, PHOX2B hindbrain, Biii Tubulin neuronal marker.

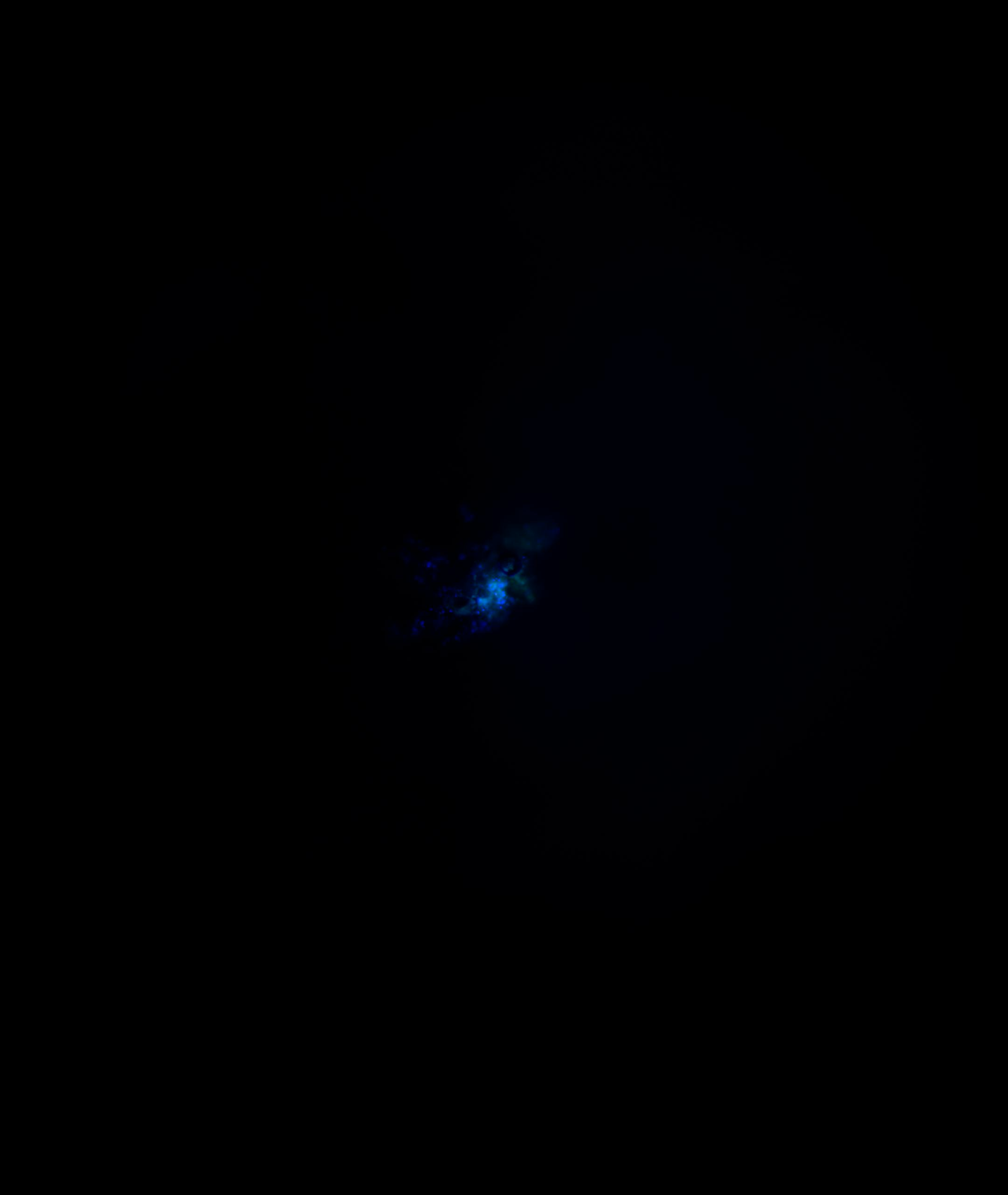
